# CTCF regulates global chromatin accessibility and transcription during rod photoreceptor development

**DOI:** 10.1101/2024.05.27.596084

**Authors:** Dahong Chen, Saumya Keremane, Silu Wang, Elissa P. Lei

## Abstract

Chromatin architecture facilitates accurate transcription at a number of loci, but it remains unclear how much chromatin architecture is involved in global transcriptional regulation. Previous work has shown that rapid depletion of the architectural protein CTCF in cell culture strongly alters chromatin organization but results in surprisingly limited gene expression changes. This discrepancy has also been observed when other architectural proteins are depleted, and one possible explanation is that full transcriptional changes are masked by cellular heterogeneity. We tested this idea by performing multi-omics analyses with sorted post-mitotic mouse rods, which undergo synchronized development, and identified CTCF-dependent regulation of global chromatin accessibility and gene expression. Depletion of CTCF leads to dysregulation of ∼20% of the entire transcriptome (>3,000 genes) and ∼41% of genome accessibility (>26,000 sites), and these regions are strongly enriched in euchromatin. Importantly, these changes are highly enriched for CTCF occupancy, suggesting direct CTCF binding and transcriptional regulation at these active loci. CTCF mainly promotes chromatin accessibility of these direct binding targets, and a large fraction of these sites correspond to promoters. At these sites, CTCF binding frequently promotes accessibility and inhibits expression, and motifs of transcription repressors are found to be significantly enriched. Our findings provide different and often opposite conclusions from previous studies, emphasizing the need to consider cell heterogeneity and cell type specificity when performing multi-omics analyses. We conclude that the architectural protein CTCF binds chromatin and regulates global chromatin accessibility and transcription during rod development.

## INTRODUCTION

Chromatin 3D organization plays a crucial role to facilitate accurate transcription at a number of loci, but how much chromatin architecture is involved in global transcriptional regulation remains elusive. Architectural protein CTCF is a central regulator of chromatin folding, and loss of CTCF leads to global dysregulation of topological domains and chromatin loops among CTCF sites^**[1, 2]**^. However, mild transcriptional changes are observed upon acute or transgenic CTCF depletion in cell culture^**[1, 2]**^ and tissues^**[3-5]**^, with limited gene expression changes in juxtaposition to CTCF binding^**[1]**^. This discrepancy is also observed when other architectural proteins are individually depleted^**[6]**^. It remains a fundamental question how broadly architectural proteins and their regulated chromatin architecture are involved in transcription regulation. A possible explanation for these results is that transcriptional changes are masked by cellular heterogeneity of tissues and cycling cell cultures. Furthermore, architectural proteins are required for normal mitosis^**[7, 8]**^, making it challenging to distinguish transcriptional effects in the context of possible mitotic arrest and apoptosis. While mitotic state and cell type resolution have not been widely considered in related studies, a few recent reports indeed revealed different regulation of chromatin architecture between synchronized and asynchronized cultured cells^**[9, 10]**^. It remains to be determined whether uniform cell populations display global transcriptional effects.

Chromatin regulators are broadly associated with developmental diseases and comprise the most abundant group among developmental disease genes, e.g., autism-risk genes^**[11]**^. While clinically attributed to neural developmental disorders^**[12, 13]**^, CTCF is widely involved in post-mitotic mouse neuronal maturation^**[3-5, 14]**^. Similar with *ex vivo* studies, CTCF depletion leads to mild gene expression changes in neural tissues^**[3-5]**^. It remains unclear whether the extraordinary heterogeneity of the nervous system masks transcriptional changes. Given the requirement for CTCF for survival of neural progenitor cells^**[15]**^, loss of CTCF may change cell composition in neural tissues, adding another layer of complexity. A major challenge is to isolate uniform cell populations and perform cell type-specific analyses in the context of a heterogenous nervous system.

Mouse retina contains post-mitotic neurons that are synchronized in a well-characterized developmental process. These uniform populations provide rare models that can help interrogate precise transcriptional regulation during development. Most mouse rods are born in a 4 d window by postnatal day 2 (P2)^**[16, 17]**^ and follow the same month-long progression, undergoing complete chromatin segregation, compaction and fusion to form single euchromatin and heterochromatin domains^**[18]**^. During this process, chromocenter numbers are reduced from ∼13 to 1, along with a nuclear inversion that places euchromatin along the nuclear periphery^**[18]**^. This extraordinary segregation and compaction reduces the nuclear granularity in rods and significantly improves vision in dim light^**[18]**^. Notably, there are various cell types with similar nuclear reorganization with undefined biological significance^**[19, 20]**^ (also see Results). Therefore, retinal cells provide excellent models of uniform post-mitotic neurons to address the relationship between transcription and chromatin architecture.

Here, we perform multi-omics analyses with sorted rods and report for the first time with *in vivo* cell type resolution that CTCF regulates global chromatin transcription and accessibility. Importantly, the transcriptomic and chromatin changes upon CTCF depletion are highly enriched for CTCF occupancy, suggesting direct CTCF regulation. CTCF mainly promotes chromatin accessibility of its direct binding targets at active chromatin but inhibits accessibility of numerous non-CTCF sites. Furthermore, CTCF binds and promotes accessibility frequently at promoters to primarily inhibit transcription. These findings provide novel and different insights from previous studies, emphasizing the importance of considering cell heterogeneity and cell type specificity when performing multi-omics analyses. We conclude that the architectural protein CTCF binds chromatin and regulates global chromatin accessibility and transcription during rod development.

## RESULTS

### CTCF primarily localizes to euchromatin

Immunochemistry typically detects broad distribution of CTCF protein throughout nuclei, for instance, in non-rod cells and newly born rods (Figure 1A, P11). During maturation, euchromatin and heterochromatin in rods segregate into peripheral and central nuclear regions ^**[18]**^, and CTCF protein becomes enriched along the nuclear periphery, where H3K4me3 euchromatin is located (Figure 1A-C, P30). This enrichment raises the question whether CTCF primarily regulates transcription in euchromatin.

**Figure 1.**
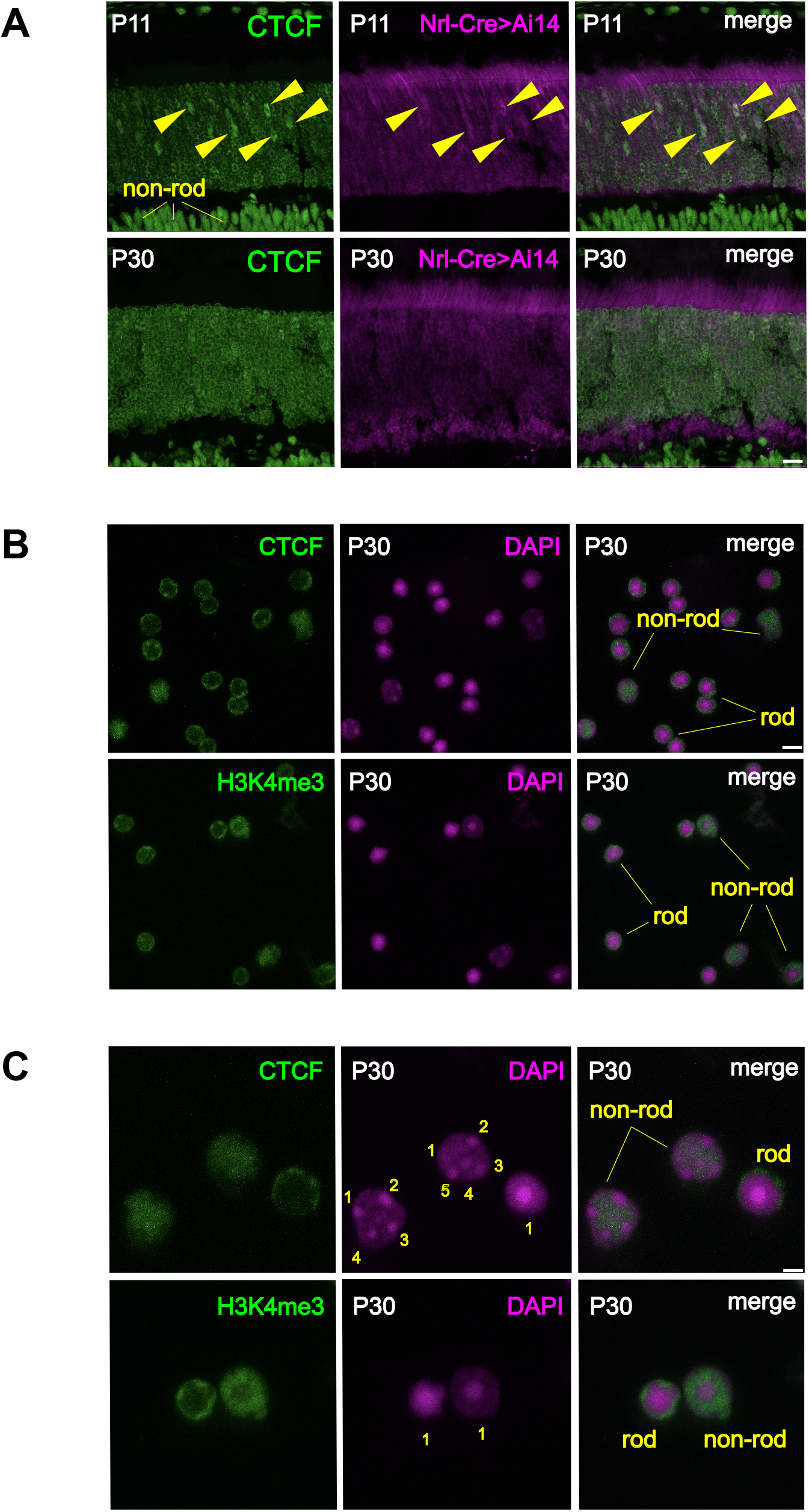
CTCF protein localizes to euchromatin. **(A)** Nrl-Cre labeled, newly born rods contain CTCF protein distributed throughout nuclei at P11 (yellow arrow heads) and becomes enriched at the nuclear periphery in developing P30 rods. In contrast, CTCF proteins are broadly distributed in non-rod nuclei throughout development. Scale bars, 10 µm. **(B)** P30 rods have small nuclei with a single central chromocenter surrounded by a thin ring of euchromatin (H3K4me3), where CTCF localizes. Scale bars, 5 µm. **(C)** Higher magnification images indicate strong segregation of chromatin in rods and different numbers of chromocenters ranging from one to multiple (numbered) among non-rods. Scale bars, 2 µm.

### Conditional CTCF depletion in rods

We crossed the well-characterized *Nrl-Cre* driver ^**[21]**^ and *CTCF*^*fl/fl***[22]**^ mice to conditionally knock out CTCF in rods. *Nrl-Cre* expression is labeled with Tomato-Ai14 and is specific for rods, with the vast majority of rods expressing strongly by P12^**[23]**^. Immunostaining indicates significantly reduced but prominent residual CTCF protein in many cKO rods at P23 (Figure 2A). CTCF protein is further diminished and minimally detectable by P30 and becomes undetectable by P35 (Figure 2A).

**Figure 2.**
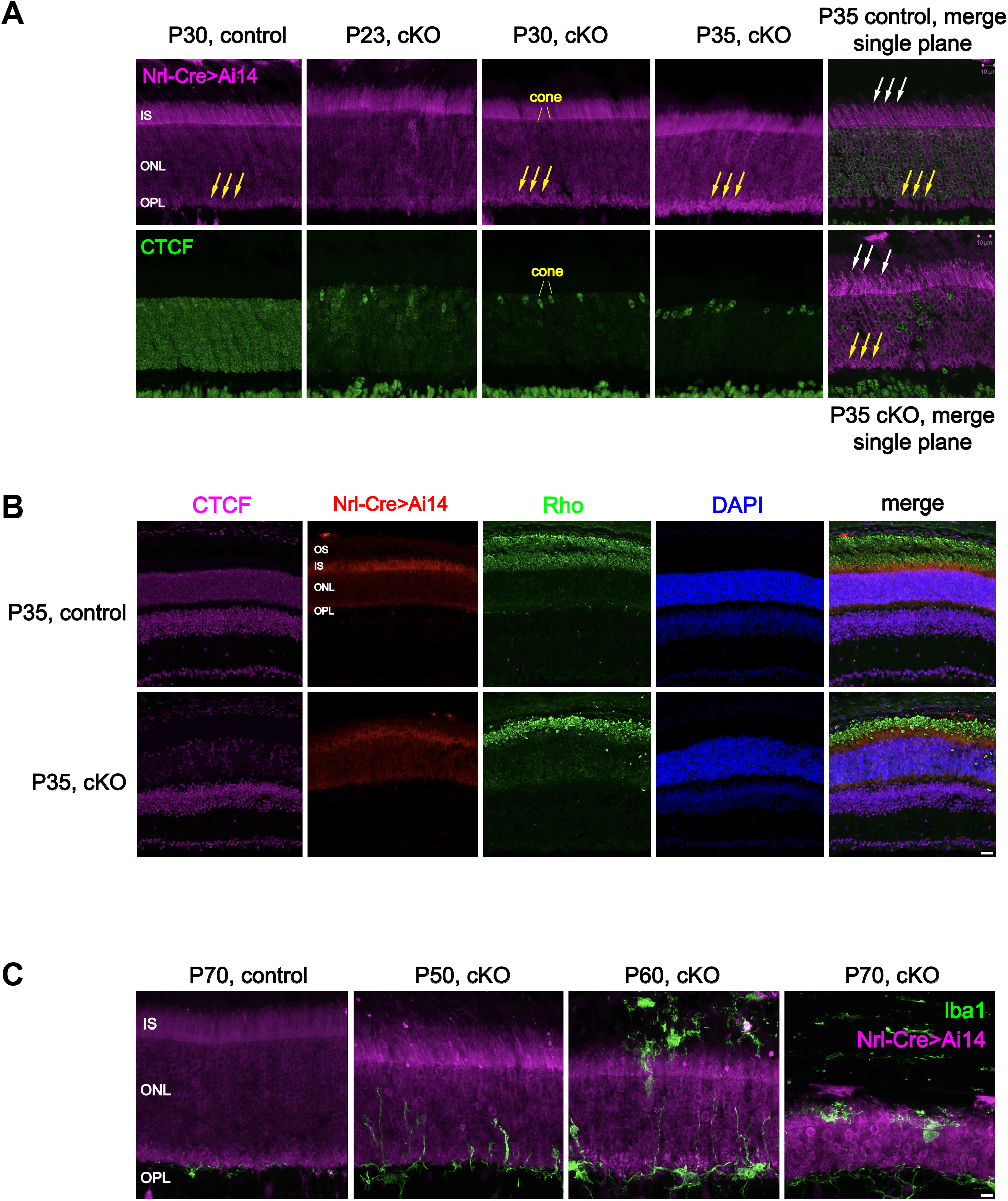
The cKO rods contain minimal CTCF proteins at P30, display morphological phenotypes at P35 and undergo degeneration in adults. **(A)** CTCF cKO rods contain reduced but prominent CTCF protein at P23, which becomes minimally detectable at P30 and undetectable at P35. CTCF protein levels stay unchanged in cones and non-rod retinal cells. Note aggregates of synaptic ribbons between photoreceptors and bipolar cells (yellow arrows), and aggregated apical inner segments (white arrows). These morphological defects become prominent at P35 and more pronounced in single confocal planes. IS, inner segment; OS, outer segments; ONL, outer nuclear layer; OPL, outer plexiform layer. Scale bars, 10 µm. **(B)** CTCF is widely depleted in rods that display aggregated outer segments at P35. Scale bars, 30 µm. **(C)** Microglia are labeled by lba1 and reside specifically in OPL of a healthy retina but upon CTCF depletion, migrate and infiltrate to clear cell debris and aggregates that eventually leads to loss of rods. Scale bars, 10 µm.

### Rod morphology and physiology are defective upon CTCF depletion

We examined morphology and physiology of CTCF-depleted rods and observed marginally detectable changes at P30. Rods reside in the outer nuclear layer (ONL) and have apical inner and outer segments (IS and OS) that contain organelles and pigment molecules, respectively. Upon light capture, rods relay signals to bipolar and amacrine cells via basal synapses at the outer plexiform layer (OPL) (Figure 2A). The cKO rods display mild aggregation of synaptic terminals at P30, represented by bigger and brighter Tomato signals in OPL (Figure 2A). At P35, both inner and outer segments, as well as synapses in OPL, display strong aggregation and prominent fragmentation (Figure 2A and B). Scavenger microglial cells, marked by Iba1 staining, are hyper-ramified and can be observed migrating from OPL into the outer nuclear layer (ONL) at P50 (Figure 2C). Microglia are fully activated at P60 and distributed throughout the ONL to scavenge rod debris. The ONL and OPL become thinner during this process due to rod degeneration through P70.

Physiological defects in cKO rods occur concurrently with morphological defects during development. Electroretinogram (ERG) measures a and b waves that indicate rod response to light stimulation and rod communication with other retinal cells, respectively^**[24]**^. The cKO mice display a trend but no statistical significance of reduced b wave amplitude compared to control at P30 (Figure 3A). By P35, the b wave becomes strengthened during normal maturation but weakened in cKO mice. This statistically significant result indicates defects in signal transduction between rods and other retinal cells, consistent with synaptic aggregation and fragmentation observed at this stage (Figure 3B). Both a and b waves are largely eliminated at P55, suggesting severe loss of photoreceptor function (Figure 3C) despite remaining rod somata. These results show that CTCF is required to ensure normal rod development and maturation both physiologically and morphologically.

**Figure 3.**
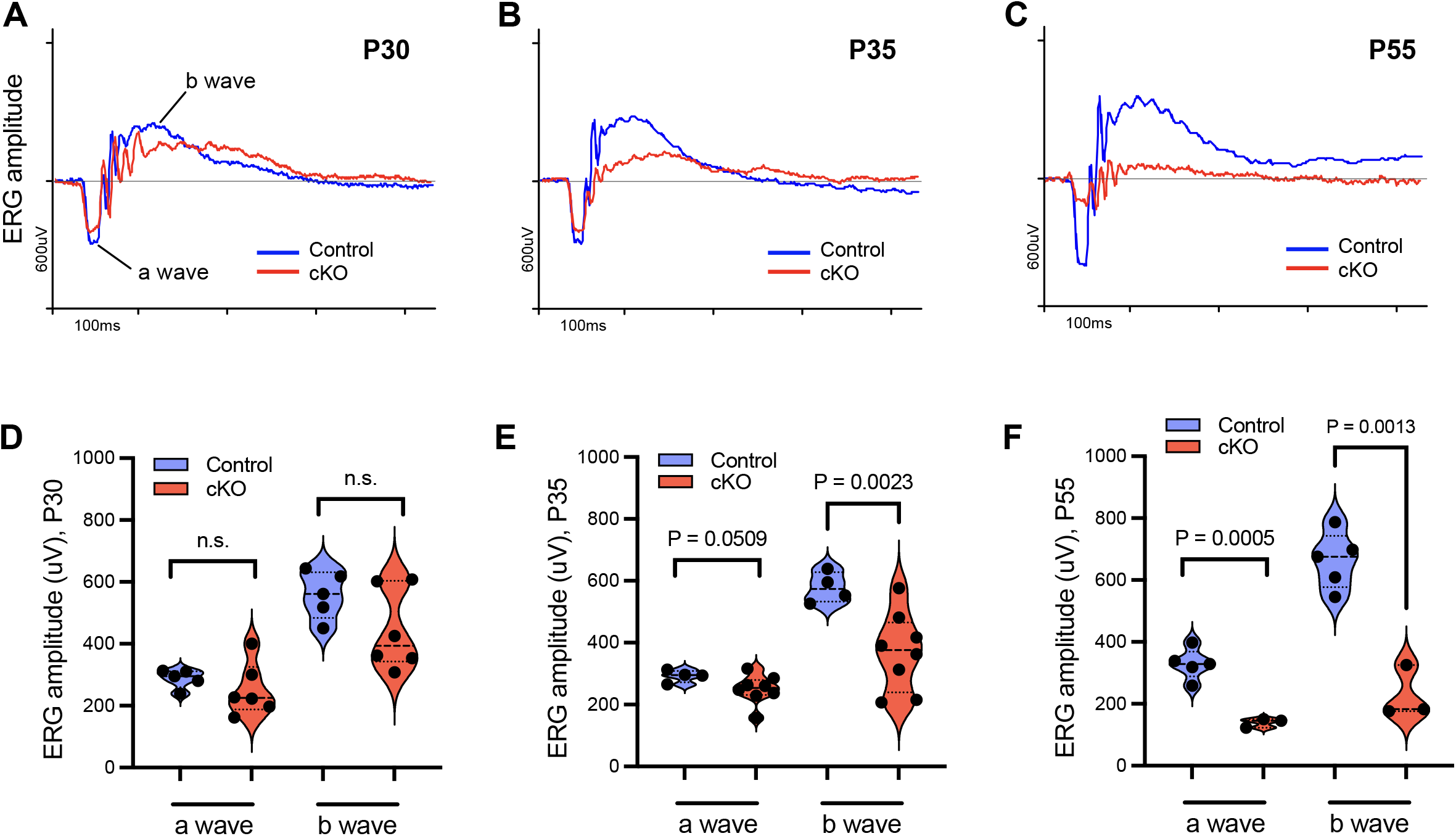
The CTCF cKO rods display physiological defects by P35. **(A-C)** ERG measures retinal response to light stimulation across a 25 d window. **(D-F)** Quantification of a and b waves indicates the first significant defects in b wave at P35. Each replicate animal is represented by a data point, and Student’s *t* test was performed to determine statistical significance.

### CTCF depletion leads to global gene expression changes

We performed RNA-seq with FACS-sorted rods and identified CTCF regulation of global gene expression. Live cKO rods were isolated at P30 when minimal residual CTCF protein can be detected but before significant phenotypes are observed, in order to minimize secondary effects. PCR duplicates were identified and removed to avoid amplification artifacts (see Methods). We identified 1,726 up-regulated and 1,541 down-regulated genes (FDR<0.05), which comprise ∼20% of the entire transcriptome (Figure 4A). We repeated the RNA-seq analyses with P35 rods and observed similar results with 497 more differential genes (Figure S1A). The mild increase of differential genes after 5 d suggests that these mutant rods exhibit limited cumulative secondary effects. Gene ontology analyses of the P30 RNA-seq data confirm strong decreased expression of *Pcdh* genes, known CTCF targets^**[5, 25]**^. Furthermore, genes involved in DNA repair, molecule binding and signal transduction are widely down-regulated (Figure 4B), consistent with the observed defects in morphology, physiology and degeneration. On the other hand, up-regulated genes are mostly enriched for those encoding zinc finger proteins, transcriptional and post-transcriptional regulators. Our analyses suggest that CTCF regulates global gene expression during rod maturation by promoting effector proteins whereas inhibiting transcriptional and post-transcriptional regulatory factors.

**Figure 4.**
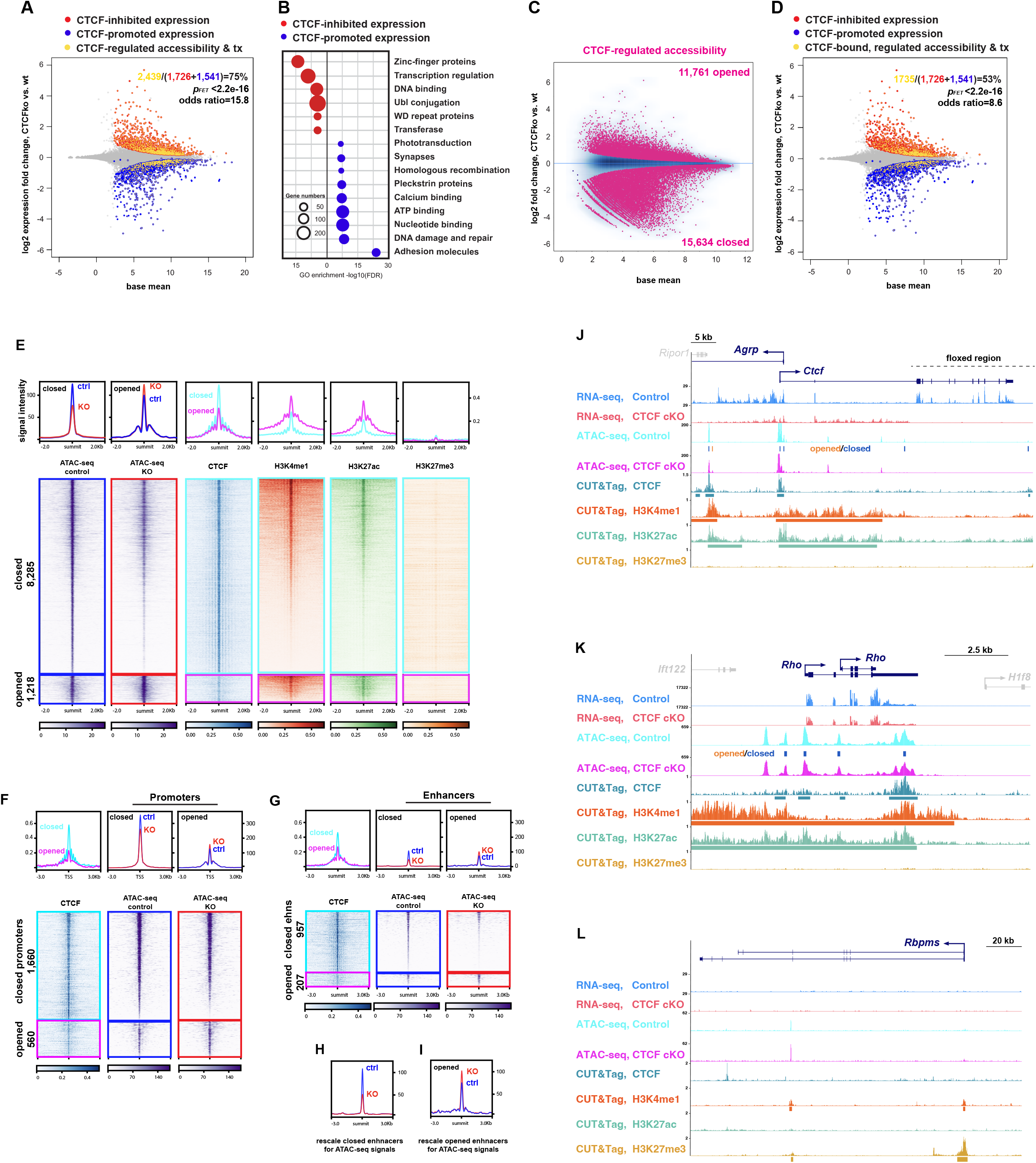
Global CTCF-regulated transcription is significantly associated with CTCF-regulated accessibility at active chromatin. **(A)** Global transcriptional changes in P30 CTCF-depleted rods are highly enriched for CTCF-regulated accessibility. FET, Fisher’s exact test; tx, transcription. **(B)** Gene ontology analyses of CTCF-regulated gene expression in P30 rods. **(C)** Global chromatin accessibility changes in P30 CTCF-depleted rods. **(D)** Loci regulated by CTCF for both transcription and accessibility are highly enriched for CTCF occupancy, suggesting direct CTCF regulation. Gene ontology analyses of CTCF-regulated gene expression. **(E)** Heatmaps of ATAC-seq signals that are regulated and occupied by CTCF in control and CTCF cKO P30 rods. Both opened and closed loci are included. CUT&Tag signals of CTCF and histone marks in control rods are also shown (one locus per row, sorted by H3K4me1 signal). Average signals across regions are color-annotated and correspond to box colors surrounding heatmaps. **(F and G)** Heatmaps and averages of ATAC-seq signals at active promoters and enhancers (H3K4me1 & H3K27ac) that are regulated and occupied by CTCF in (E). **(H and I)** Rescales of averaged ATAC-seq signals at enhancers in (G). Note the stronger accessibility changes at enhancers than at promoters (F). **(J)** The floxed *Ctcf* locus indicates loss of transcription of all CTCF-coding sequences and the reduced expression of a juxtaposed gene upon CTCF depletion. **(K and L)** Cell type specificity of FACS purification is confirmed by high expression of the rod-specific gene *Rho* and no expression of the retinal ganglion cell-specific gene *Rbpms* in the RNA-seq data. This transcriptional specificity is also reflected by the active and inactive histone marks at *Rho* and *Rbpms*, respectively.

### CTCF depletion disturbs global chromatin accessibility that is enriched at differentially expressed genes

Our previous study indicates that proper chromatin accessibility is crucial for temporal chromatin architecture and transcription at specific loci during neuronal development^**[26]**^. We therefore tested whether CTCF regulates global chromatin accessibility by performing ATAC-seq with sorted live P30 rods. We identified 27,395 of 66,520 ATAC-seq peaks with differential accessibility upon CTCF depletion, which comprise >41% of genome-wide accessibility (Figure 4C). These differentially accessible loci contain more closing genomic loci (15,634) with stronger accessibility changes than opening loci (11,761) (Figure 4C). Remarkably, 75% (2,439 of 3,267) of the CTCF-regulated differential genes are also regulated by CTCF for accessibility (Figure 4A). This strong enrichment (*p*<2.2e-16, odds ratio=15.8, Fisher’s Exact Test (FET)) suggests that CTCF-dependent chromatin accessibility plays a key role in CTCF-mediated transcriptional regulation.

### CTCF directly binds and regulates chromatin accessibility at differential genes

In order to assess direct CTCF regulation, we performed CUT&Tag of CTCF with sorted P30 control rods and observed enrichment of CTCF binding at loci regulated by CTCF for both chromatin accessibility and transcription. We identified significant enrichment of CTCF occupancy (FET *p*<2.2e-16, odds ratio=2.9) at 9,503 of 27,395 (35%) loci that harbor CTCF-dependent accessibility. More than 87% (8,285 of 9,503) of these CTCF targets become closed upon CTCF depletion (Figure 4E), suggesting that CTCF binding promotes chromatin accessibility in rods. This result differs from the comparable numbers of closed and opened CTCF binding targets upon CTCF depletion in a previous study^**[27]**^.

We further integrated RNA-seq, ATAC-seq and CUT&Tag to determine whether CTCF binding and accessibility regulation are associated with transcriptional consequences. About 60% (1,949 of 3,267) of differential genes are direct CTCF binding targets, of which 72% (1,735 of 1,949) are also regulated by CTCF for chromatin accessibility, displaying strong enrichment of CTCF binding among its accessibility and transcriptional targets (FET *p*<2.2e-16, odds ratio=8.6) (Figure 4D). Therefore, our data suggests global, direct CTCF binding and regulation of chromatin accessibility that are associated with transcriptional responses.

### Strong CTCF-binding promotes accessibility of active chromatin

To determine what type(s) of chromatin are bound and regulated by CTCF, we performed CUT&Tag of histone modifications with sorted P30 control rods. Among the ATAC-seq loci bound and regulated by CTCF, 48% (4,568 of 9,503) are covered and significantly enriched (FET *p*<2.2e-16, odds ratio=2.3) for H3K4me1 (Figure 4E), which labels bivalent proximal promoters and distal enhancers^**[28-30]**^. We found that 10% (457 of 4,568) of these loci are also labeled by the H3K27me3 mark, suggesting a poised state. Moreover, 74% (3,384 of 4,568) harbor the active mark H3K27ac (FET *p*<2.2e-16, odds ratio=22.3), suggesting enriched CTCF regulation at active chromatin. This result is consistent with CTCF subnuclear distribution overlapping euchromatin (Figure 1). Approximately 74% (2,617 of 3,384) of these CTCF-regulated active elements are closed upon CTCF depletion (Figure 4E), suggesting that promoting accessibility is a major function of CTCF binding at active chromatin.

Regarding genomic location, two thirds of all CTCF-regulated active loci (2,220 of 3,384) are located within ±1.5 kb of transcription start sites (Figure 4F), indicating preferential CTCF regulation of proximal active promoters relative to distal active enhancers (Figure 4F and G). Notably, accessibility changes are stronger in amplitude at active enhancers than promoters (Figure 4F-I). Consistent with these results, enhancers were recently identified as major targets for accessibility regulation during post-mitotic neuronal development^**[26, 31, 32]**^.

Aside from H3K4me1-marked regions, there are 4,935 ATAC-seq regions occupied and regulated by CTCF that harbor moderate H3K27ac (560) or H3K27me3 (737) (Figure S1B). Almost all of these regions (94%; 4,641 of 4,935) are closed upon CTCF depletion, confirming global CTCF binding-mediated promotion of chromatin accessibility. Our findings contrast strongly with previous studies, which showed low CTCF occupancy at proximal regions^**[2, 33]**^ and suggested primary CTCF function in repressing chromatin accessibility^**[27]**^.

Remarkably, CTCF-binding strength is stronger at CTCF-promoted loci than CTCF-inhibited loci, regardless of chromatin state or genomic location (Figures 4E-I and S1B), implying a positive correlation between CTCF binding strength and CTCF function in promoting accessibility in rods. This correlation is different from a previous report suggesting that strong CTCF binding inhibits chromatin accessibility^**[27]**^.

### CTCF binding promotes accessibility and results in more transcriptional repression than activation

We noticed the discrepancy that there are more up-regulated than down-regulated genes (758 up vs. 637 down) compared to the large majority of closed CTCF targets upon CTCF depletion (8,285 closed vs. 1,218 opened) (Figure 4E). By focusing on closed CTCF targets, we again identified more up-regulated genes (669 up vs. 619 down) in juxtaposition (Figure 5A), although some genes overlap both closed and opened regions. We surprisingly noticed 220 up-regulated genes (Figure 5B) that contain solely closed chromatin corresponding to 280 regions. Motif analyses of these 280 closed regions indicate the highest motif enrichment of CTCF, as expected, followed by CTCFL, which is however not expressed in rods (Figure 5C and S2B). Binding motifs are also significantly enriched for the Sp/KLF family and PATZ1, both of which may act as transcription repressors. The Sp/KLF family contains many zinc finger proteins that compete to repress or activate transcription in a promoter and cell type-specific manner^**[34, 35]**^. PATZ1 is a chromatin remodeling factor and putative insulator factor that represses transcription^**[36-38]**^. Loss of CTCF reduces chromatin accessibility, which may lower occupancy of these transcription repressors and result in ectopic transcription. We also performed motif enrichment analyses with all the 8,285 closed CTCF binding targets and identified identical motifs with stronger statistical significance due to more genomic regions being analyzed (Figure 5C). Our results suggest that these transcription repressors might cooperate with CTCF globally to regulate transcription.

**Figure 5.**
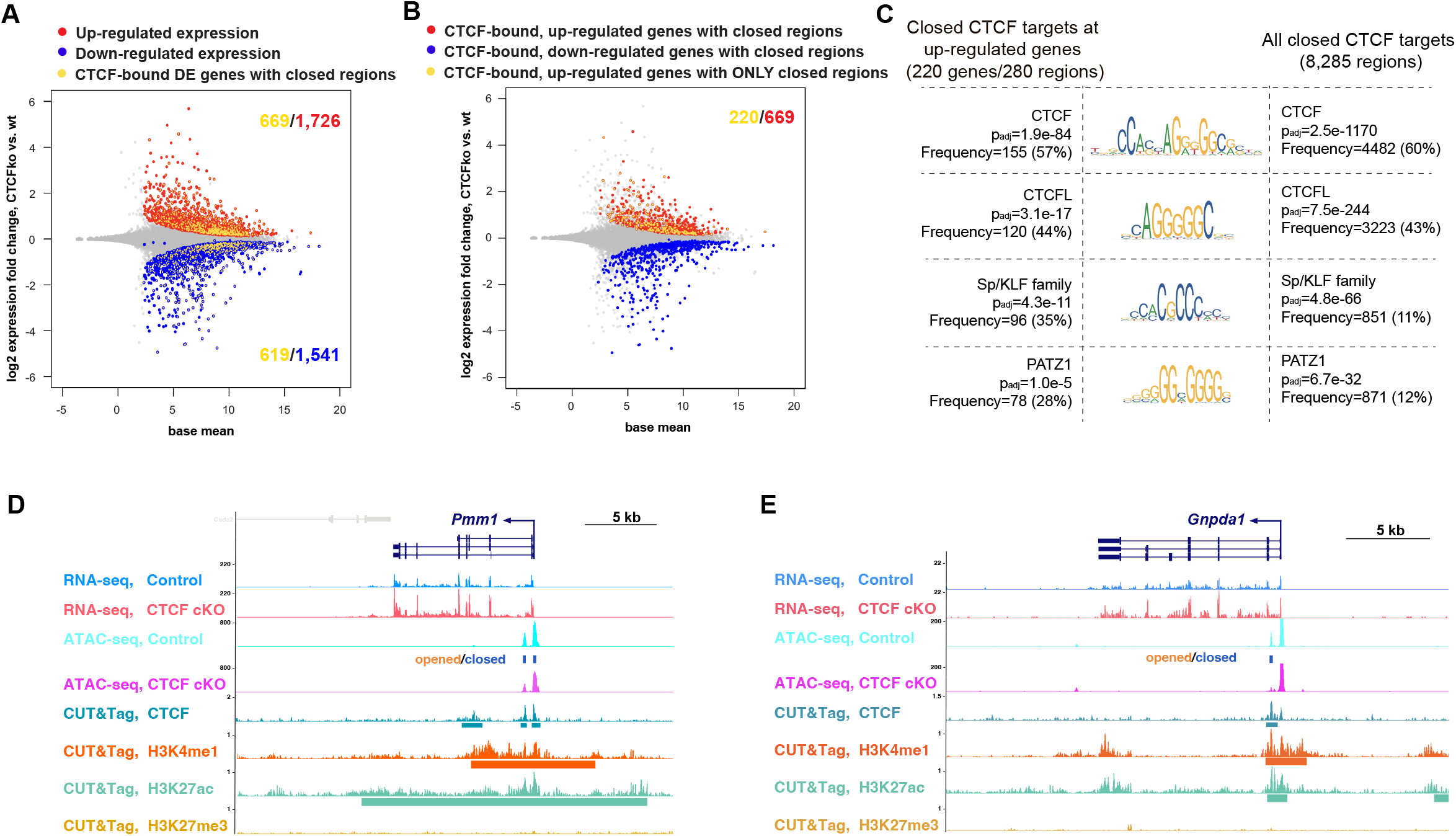
CTCF promotion of chromatin accessibility corresponds to substantial transcription inhibition. **(A)** CTCF promotion of chromatin accessibility leads to both promotion and inhibition of expression. **(B)** There are 220 solely closed genes with higher expression upon CTCF depletion. **(C)** Both the 220 genes and global targets of CTCF-inhibited accessibility are enriched with binding motifs of transcription repressors identified by motif enrichment analyses. **(D and E)** Representative loci that contain CTCF-promoted accessibility at active chromatin with CTCF-inhibited transcription.

Aside from the closed regions, 1,218 opened CTCF-binding targets include 252 up-regulated and 67 down-regulated genes in juxtaposition, consistent with the typical trend in which chromatin accessibility positively correlates with transcription. Motif enrichment analyses of these 1,218 regions identify again identical motifs as the closed regions for CTCF that identify the Sp/KLF family and PATZ1, suggesting that interactions and likely competition among these transcription regulators are globally applicable and lead to diverse transcriptional consequences.

### CTCF remotely inhibits chromatin accessibility

Beyond the 9,503 CTCF-bound targets, there are 17,847 genomic regions that harbor CTCF-dependent accessibility but not CTCF occupancy. In contrast to CTCF-occupied regions, there are more opened than closed regions among these non-CTCF regions (10,498 opened vs. 7349 closed) (Figure S2). These non-CTCF regions have a similar composition to CTCF targets, consisting of 40% promoters (7,098 of 17,847) and 31% (5,449 of 17,847) H3K4me1-labeled distal enhancers, most of which are active enhancers (3,075 of 5,449) with minimal poised enhancers (443 of 5,449) (Figure S2). We speculate that some opened regions may be located at Polycomb domains and become activated by spreading of active transcription due to loss of CTCF and insulator activities. However, only a small proportion of these opened promoters (802 of 4,787) and enhancers (26 of 2,253) are covered by H3K27me3 in control P30 rods, suggesting that their accessibility and transcription are unlikely regulated by Polycomb repressive complexes. Remarkably, the inhibition of accessibility at these non-CTCF regions confirms the correlation between CTCF function and binding strength (Figure S2), and we speculate that CTCF regulates accessibility of these remote loci via long-distance loops.

## DISCUSSION

By performing cell type-specific multi-omics analyses, we elucidated CTCF regulation of global chromatin transcription and accessibility during rod development. The global, CTCF-dependent transcriptomic and epigenomic changes are highly enriched for CTCF occupancy, indicating direct and enriched CTCF regulation of active chromatin. We further elucidated dimorphic CTCF functions in promoting and repressing chromatin accessibility that strongly correlate with CTCF binding strength. Finally, our findings reveal novel factors whose binding motifs overlap with CTCF occupancy in rods to promote chromatin accessibility while repressing transcription. To our knowledge, this is the first report of CTCF chromatin-related functions during post-mitotic development with cell type resolution. Our findings provide novel insights into relationships among architectural proteins, transcriptome and epigenome. Our study strongly suggests that cell heterogeneity, mitotic state and developmental context are all worth careful consideration during multi-omics analyses.

### CTCF regulates global transcription

Depletion of individual architectural proteins such as CTCF, Rad21, WAPL, or NIPBL leads to dysregulation of global chromatin architecture but mild gene expression changes. This discrepancy has been confirmed in studies utilizing cell culture and tissues coupled with acute protein degradation or conditional knockout, but the underlying reasons and mechanisms remain unclear. Notably, AID tagging strongly lowers endogenous CTCF expression before auxin treatment^**[1, 27]**^, thus RNA-seq analyses may only reveal mild differences between hypomorphic and null cells.

We speculate that cellular heterogeneity in previous studies also masks transcriptional changes from being fully determined. Our transcriptomic analyses with sorted rods detect >3,000 differential genes. UMI-guided PCR deduplication further improves quantification accuracy. We did not analyze rods earlier than P30 due to prominent residual CTCF protein (Figure 2A), which is likely to be functional below 90% depletion^**[1]**^. To evaluate progressive secondary effects, we observed slightly higher (∼400) differential genes at P35 than P30, which are an order lower than previous studies. The fact that these knockout rods survive more than two months also implies that cumulative secondary changes occur very slowly.

More importantly, our observed global transcriptional changes are highly enriched for CTCF binding and CTCF-regulated accessibility, suggesting direct CTCF regulation. We detected CTCF binding at >60% of differential genes, slightly higher than a previous study (∼50%), which performed acute CTCF degradation in mouse ES cells^**[1]**^. Furthermore, ∼43% of differential genes contain CTCF binding and CTCF-dependent accessibility in our analyses, while ∼16% differential genes contain CTCF binding in another recent study of acute CTCF degradation in a human B-cell line^**[27]**^. Taking all evidence into consideration, we believe the global transcriptional changes upon CTCF depletion mostly result from loss of CTCF, and it is essential to consider cellular heterogeneity when performing global multi-omics studies.

### CTCF promotes accessibility of active chromatin in rods

Our study indicates that CTCF promotes accessibility of its binding targets, which are highly enriched for euchromatin at active promoters and enhancers. This observation appears different from the previously described comparable closing and opening upon CTCF depletion in cell culture^**[27]**^. Furthermore, we observed tight correlation between CTCF promotion of chromatin accessibility and strong CTCF binding, which inhibited accessibility in a previous study^**[27]**^. We speculate that the differences result from distinctive cell types (rods vs. leukemia cell culture), cycling states (post-mitotic vs. mitotic), and/or cellular environments (*in vivo* vs. *ex vivo*).

### CTCF promotes accessibility and represses transcription

We identified more up-regulated than down-regulated genes, despite the predominant closure of active chromatin upon CTCF depletion. There are 220 up-regulated genes that harbor only closed chromatin at promoters and active enhancers, which are enriched for binding motifs of transcription repressors (Figure 5). The closed chromatin may expel transcription repressors to allow for increased transcription. An alternative but not mutually exclusive possibility is that the closed chromatin harbors silencers that interact with target genes less frequently due to reduced accessibility, which results in increased transcription.

### Rods as a model to study transcription and nuclear architecture

Rods undergo extensive chromatin segregation and form single euchromatin and heterochromatin domains that surround a single chromocenter. This extensive segregation leads to low nuclear granularity that reflects particular cellular functions and states, i.e., light detection in rods. Notably, similar nuclear organization is also employed by other types of cells, e.g., olfactory neurons undergo similar chromatin segregation and nuclear inversion to achieve low nuclear granularity^**[19]**^. Non-rod retinal cells also contain a wide range of nuclear organization and granularity^**[39]**^, harboring 1-5 chromocenters of various sizes (Figure 1B and C). Immune cells are well studied for their diverse nuclear organization and granularity, which reflects distinct functions and differentiation states^**[40]**^. Whether cells with high levels of chromatin segregation and low nuclear granularity share similar architectural protein functions with rods to regulate global transcription remains unknown. Studying how cells with different levels of chromatin separation and nuclear granularity respond to loss of architectural proteins will help elucidate relationships between transcription and different levels of chromatin architecture, such as chromatin folding and compartmentalization. Given the minimal overlap between CTCF proteins and high-DAPI heterochromatin regions regardless of cell types (Figure 1C), CTCF function and mechanisms learned in rods may be shared by other cell types as well. We believe that rods provide an excellent opportunity to study how transcriptome, epigenome and chromatin architecture interact in a context of high chromatin segregation.

## METHODS

### Animals

Founder mice were genotyped to exclude mutations *rd*^*1*^, *rd*^*8*^ and *rd*^*10*^ and backcrossed to C57BL/6J mice for eight generations. CTCF conditional knockout in rods was achieved by crossing previously described mice that carry *Nrl-Cre*^**[21]**^, *CTCF*^*fl/fl***[22]**^ and *Tomato-Ai14*^**[23]**^. Age-matched mice with Cre and Tomato were used as controls. Both sexes were used for experiments except for multi-omics assays, which used only males to assess both X and Y chromosomes. Rods were sorted either in late afternoon of P30 (RNA-seq and ATAC-seq) or early morning of P31 (CUT&Tag) and are collectively presented as P30 rods. Control and cKO samples have eight and three biological replicates, respectively.

### Fluorescence-activated cell sorting (FACS)

Retinas were retrieved, dissociated and FACS-sorted following a published protocol^**[41]**^. Nuclei were labeled using Ruby (Thermo Fisher Scientific V10309), and dissociated cells were selected against dead and dying cells using DAPI. Rods were selected as cells with Ai14 Tomato labeling and low granularity^**[39]**^ on a FACSAria II machine in the Flow Cytometry Core at the National Heart, Lung and Blood Institute. Cells were either frozen in lysis buffer for RNA-seq or used for downstream applications immediately after sorting.

### Immunostaining, imaging and quantification

Isolated retinas were crosslinked for 1 h in 4% PFA and soaked in 15% and 30% sucrose over two nights. Retinas were then OCT-embedded and cryo-sliced into 16 μm sections. Immunostaining was performed following a published protocol^**[18]**^ with primary antibodies at 1:1,000 to visualize CTCF (Cell Signaling, D31H2), Rhodopsin (Abcam, ab5417), H3K4me3 (Abcam ab8580), and Iba1 (FUJIFILM, 019-19741). Secondary antibodies (Thermo Fisher Scientific) were used at 1:1,000. Images were taken as maximum-intensity z-series projections with a Zeiss 780 at NIH or a Leica SP8 confocal microscope at University at Buffalo.

### RNA-seq libraries, sequencing and analysis

RNA was extracted from sorted rods with RNeasy Plus Micro Kit (Qiagen) according to manufacturer’s protocol and was quantified with Quant-iT RiboGreen RNA Assay Kit (Thermo Fisher Scientific). RNA integrity was confirmed with High Sensitivity RNA ScreenTape, and 1 ng total RNA was used to generate RNA-seq libraries with SMARTer Stranded Total RNA-Seq Kit v3 (Takara). All samples were sequenced with NovaSeq 6000 (Illumina) at the NHLBI Genomics Core Facility by 50 bp paired-end sequencing.

We ran UMI-tools to identify unique molecular identifiers (UMI), removed PCR duplicates and retained only reads that represent individual RNA molecules for quantification. Using cutadapt v1.10, we performed quality trimming, and poly A and adapter trimming. To identify differentially expressed genes, the deduplicated and trimmed reads were aligned with hisat2 v2.0.4 to the GENCODE Release M31 mouse assembly. Reads were counted for exons of annotated genes using featureCounts (v1.5.0-p3) in paired-end mode. Counts were loaded into R, and the DESeq2 package was used to identify differentially expressed genes (FDR<0.05) with a read count threshold at 58. Counts from all libraries were loaded together for normalization and size factor calculation, which was further used to visualize data in UCSC. Similar results of GO analysis were obtained with PANTHER^**[42]**^, GOrilla^**[43]**^ and DAVID^**[44]**^. DAVID results with FDR < 1e-4 were used to generate the plot for CTCF-regulated genes. Redundant GO terms were presented only once in each category.

### ATAC-seq libraries and analyses

Omni-ATAC-seq libraries were generated according to a detailed protocol^**[45]**^ with minor adjustments. Specifically, 3 × 10^5^ sorted rods were used for each biological replicate. Eight total PCR cycles were performed to amplify libraries, which were subsequently double size-selected with AMPure XP beads (first with 0.6x volume, then with 1.2x volume) (Beckman Coulter). Finally, 50 bp paired-end sequencing was performed at the NHLBI Genomics Core Facility on an Illumina NovaSeq 6000.

Adapter sequences were trimmed from reads with cutadapt (v2.3) and aligned to the GENCODE Release M31 mouse assembly with bowtie2 (v2.3.5; --very-sensitive, paired-end mode). Reads were depleted for mitochondria alignment and for multi-mapped reads with samtools (v1.9). Uniquely mapped reads were further depleted for PCR duplicates with picard and computationally size-selected for inserts <150 bp for reads from nucleosome-free regions. ATAC-seq peak calling was performed with MACS2 (v2.2.6; pair-end mode -f BAMPE), and differential accessibility was called with the R package DiffBind v3 with default parameters. Motif enrichment analyses were performed with AME 5.3.336 and SEA 5.5.5, and only motifs identified by both algorithms with known binding factors were reported.

### CUT&Tag libraries and analyses

After FACS sorting, collected rods were used directly to generate CUT&Tag libraries following a published protocol (https://dx.doi.org/10.17504/protocols.io.bcuhiwt6) with minor changes. Specifically, 1.4 × 10^5^ rods were used for each biological replicate without crosslinking. Tn5 was purchased from Epicypher (15-1017) and used at 1:20. The primary antibodies CTCF (Cell Signaling Technology D31H2), H3K27me3 (Cell Signaling Technology 9733), H3K4me1 (Abcam ab8895) and H3K27ac (Abcam ab4729) were used at 1:50. Fourteen total PCR cycles were performed to amplify libraries, which were re-purified (using 1.1x volume of AMPure XP beads) after pooling to remove primer dimers. Then 50 bp paired-end sequencing was completed at the NHLBI Genomics Core Facility on a NovaSeq 6000. Computational analyses of CUT&Tag data were performed with the same pipeline used for ATAC-seq except that there was no computational size-selection of sequencing reads.

### Electroretinogram Recording

All procedures were performed in a dark room with dim red light. Mice were dark-adapted overnight and anesthetized with ketamine (100 mg/kg)/xylazine (6 mg/kg) mixture. Mice were placed on a 37°C heating pad, and 1% tropicamide and 0.5% phenylephrine were applied to eyes. ERG was performed using an Espion E2 system (Diagnosys) that was connected to a LED colordome. Responses were recorded using gold loop electrodes in contact with the cornea, plus a mouth reference electrode and a tail ground electrode. ERG was captured after a flash (<4 ms) at 10 cd·s/m^2^. Statistical analyses of ERG amplitudes were performed with Student’s *t* test in Prism 10, and *p* values and data dots were labeled in violin plots.

### Statistics

Fisher’s Exact Test analyses were performed in R, and *p* values beyond the machine epsilon were returned as zero in R and reported as <2.2e-16 in this paper.

## Acknowledgements

We thank Shiming Chen at WUSTL for sharing animals; Haohua Qian and Yichao Li at NEI for assistance with ERG training and discussion; Douglas Forrest, Lily Ng and Mallika Bhattacharya at NIDDK for comments on the results. This work was funded by the Intramural Program of the National Institute of Diabetes and Digestive and Kidney Diseases, National Institutes of Health (DK015602 to E.P.L.) and the NIH Pathway to Independence Award (K99HD097308-01A1 and 4R00HD097308-02 to D.C.).

## Author contributions

Conceptualization: D.C.; Data Curation: D.C.; Formal Analysis: D.C., S.W.; Investigation: D.C.; Methodology: D.C., S.K.; Resources: D.C., E.P.L.; Software: D.C., S.W.; Validation: D.C.; Visualization: D.C., S.W.; Writing – original draft: D.C.; Writing – review & editing: D.C., S.K., S.W., E.P.L.

## LEGENDS

**Figure S1.**
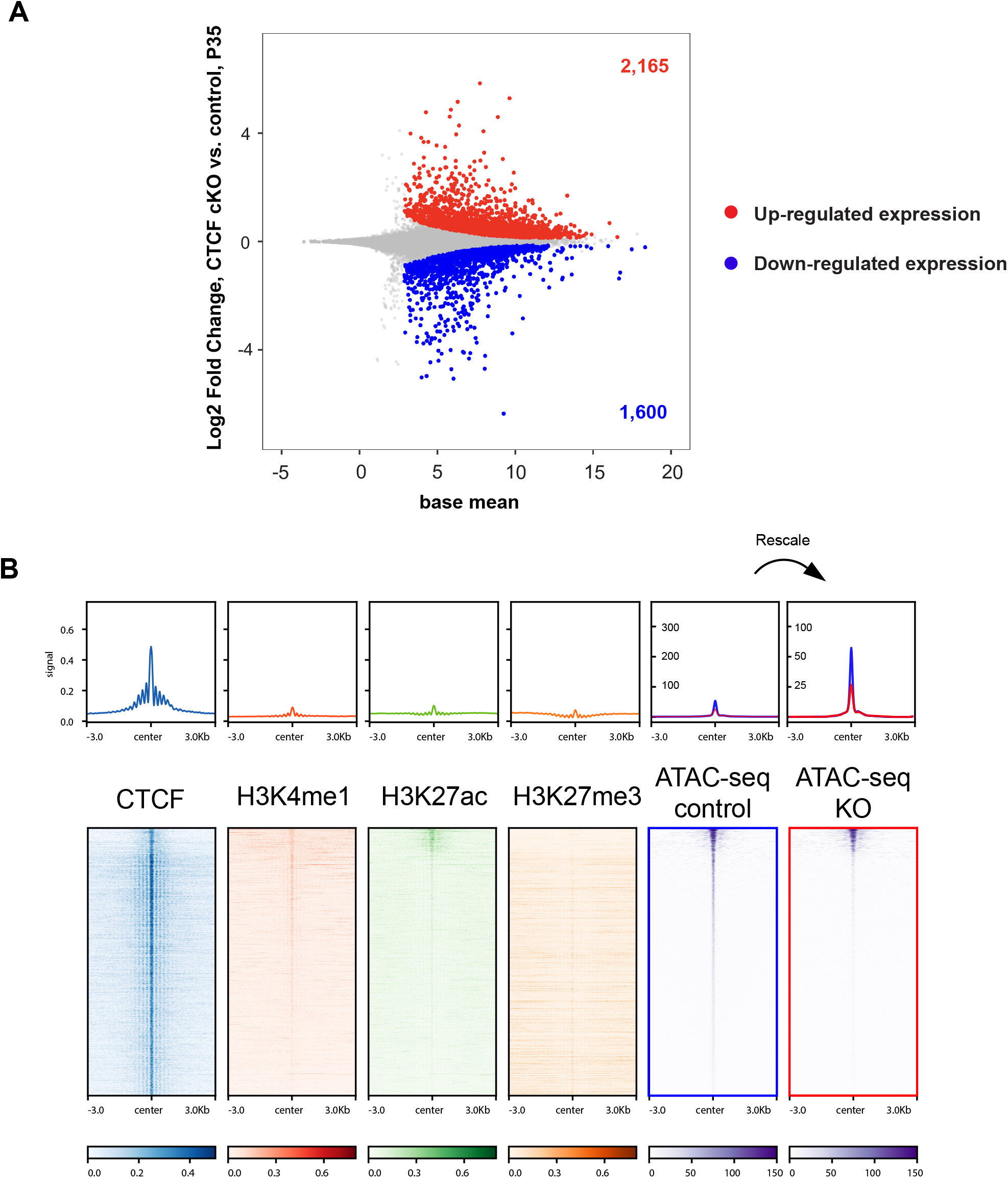
CTCF-regulated gene expression in P35 rods and CTCF-regulated chromatin accessibility at non-histone mark loci in P30 rods. **(A)** Loss of CTCF leads 2,165 up-regulated and 1,600 down-regulated genes in P35 rods at *p*_*adj*_ < 0.05. **(B)** Heatmaps and averaged signals are shown for CTCF-bound and -regulated loci that contain no histone marks. Note the strong CTCF binding and closing at the vast majority of regions in CTCF cKO rods. Rescale of the averaged accessibility signals are also shown.

**Figure S2.**
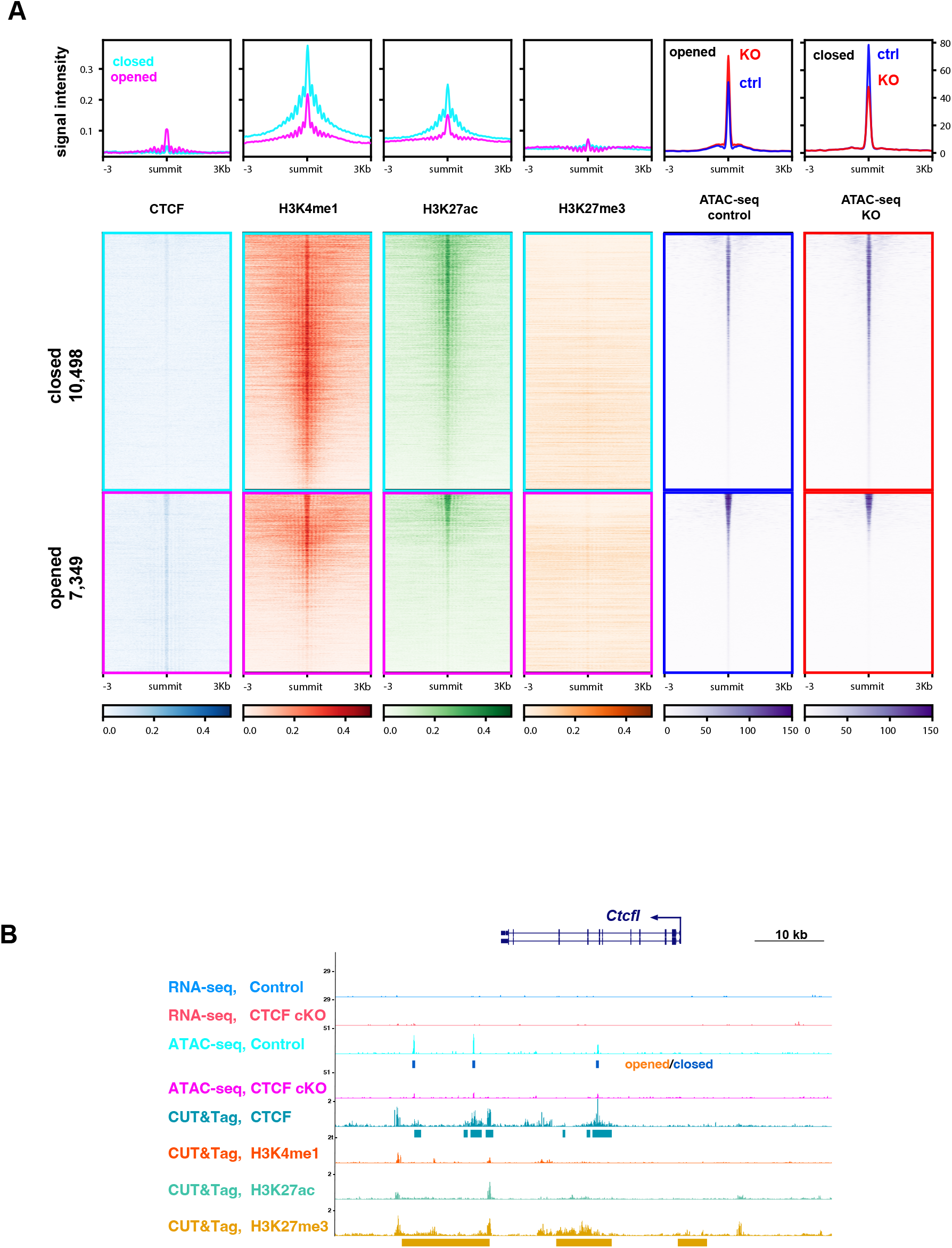
Heatmaps of accessibility changes at non-CTCF loci. **(A)** There are slightly more opened than closed regions upon CTCF depletion at regions that are not occupied by CTCF. **(B)** *Ctcfl* is not transcribed in rods.

